# Decoding the auditory brain with canonical component analysis

**DOI:** 10.1101/217281

**Authors:** Alain de Cheveigné, Daniel Wong, Giovanni M. Di Liberto, Jens Hjortkjaer, Malcolm Slaney, Edmund Lalor

## Abstract

The relation between a stimulus and the evoked brain response can shed light on perceptual processes within the brain. Signals derived from this relation can also be harnessed to control external devices for Brain Computer Interface (BCI) applications. While the classic event-related potential (ERP) is appropriate for isolated stimuli, more sophisticated “decoding” strategies are needed to address continuous stimuli such as speech, music or environmental sounds. Here we describe an approach based on Canonical Correlation Analysis (CCA) that finds the optimal transform to apply to both the stimulus and the response to reveal correlations between the two. Compared to prior methods based on forward or backward models for stimulus-response mapping, CCA finds significantly higher correlation scores, thus providing increased sensitivity to relatively small effects, and supports classifier schemes that yield higher classification scores. CCA strips the brain response of variance unrelated to the stimulus, and the stimulus representation of variance that does not affect the response, and thus improves observations of the relation between stimulus and response.

## 1 Introduction

Common techniques to measure brain activity include electroencephalography (EEG), magnetoencephalography (MEG), electrocorticography (ECoG), and functional magnetic resonance imaging (fMRI). Paired with controlled sensory stimulation, such techniques allow sensory and perceptual processes to be probed. In the case of EEG and MEG, one common approach has been to examine how the time course of brain potentials/fields is affected by particular stimuli. However, stimuli usually need to be repeated several times, and the recorded signals averaged to overcome the many sources of noise and brain activity unrelated to the stimulation. This is practical only for short stimuli or isolated events, and precludes the study of responses to longer and more naturalistic stimuli such as speech, music, or environmental sound. Recently, new approaches based on system identification allow meaningful response functions to be derived from ongoing stimulation.

A short event such as a click or sound onset produces a stereotyped EEG/MEG response, with peaks and troughs that are referred to by conventional names (N100, P300, etc.). Their morphology, timing and dependency on experimental conditions have yielded a wealth of information about perceptual processes (Woodman, 2010). Activity related to ongoing stimulation is harder to interpret because responses to each individual event overlap in time, and the lack of repetition precludes simple averaging over trials. Nonetheless, supposing that the system is *linear* and *time-invariant,* the relation between stimulus and response can be described as a convolution, and characterized by an impulse response or *temporal response function* (TRF) that can be estimated from the stimulus and response pair by systems identification techniques (Lalor et al., 2009; Crosse et al., 2016).

This linear model can be estimated and evaluated in either of two ways. In the first (“forward model”) the model is used to predict the neural response from the stimulus, and the prediction is compared to the actual measurement. In the second (“backward model”, or “stimulus reconstruction”), the model is used to infer the stimulus from the response, and the inferred stimulus is compared to the actual stimulus. The quality of fit may be quantified in terms of correlation coefficient, and results cross-validated by measuring the correlation on data distinct from the data that served to train the model. The model can also be used to design a classifier, and its quality quantified by percentage correct classification, or area under the Receiver Operating Curve (ROC).

With either approach correlation scores are usually modest (on the order of r=0.1-0.2 for EEG data) albeit statistically significant. Such low scores are sobering, but they should not come as a surprise. EEG signals reflect many brain processes in addition to those related to sensory processing, and thus only a fraction of EEG variance can be predicted from the stimulus (Woodman, 2010). Conversely, certain features of the stimulus may have little or no impact on the percept or brain activity evoked. Low correlation scores thus reflect the fact that EEG and stimulus each include variance that is irrelevant to the perceptual process.

In this paper we explore a third approach: transform both stimulus and response so as to minimize irrelevant variance, and evaluate the quality of the model by measuring the correlation after each data set has been transformed. This allows the stimulus representation to be stripped of dimensions irrelevant for measurable brain responses, and the EEG to be stripped of activity unrelated to auditory perception. The method we use is based on *canonical correlation analysis* (CCA) (Hotelling, 1936). Given two sets of data (here stimulus and EEG), CCA finds the best linear transform **W_1_** to apply to the first to maximize its projection on the second, and the best linear transform **W_2_** to apply to the second to maximize its projection on the first. CCA is a linear technique, but it can be extended to characterize non-linear and convolutional relations by including appropriate transforms of the data before applying CCA (see Discussion).

## 2 Methods

### EEG/MEG data model

Electrical activity within the brain is picked up by electrodes on the skull (EEG) or magnetometers (MEG) surrounding the head. The source-to-sensor relation is usually linear: the *J* measured brain signals *b_j_* (*t*) at *T* time steps, forming a matrix of dimensions *T* × *J*, are linearly related to the activities of *I* sources *s_i_*(*t*) within the brain:

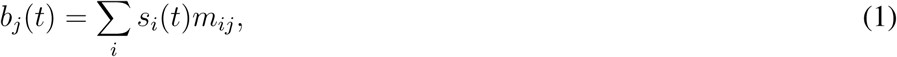

where *t* is time and the *m_ij_* are unknown source-to-sensor mixing weights. In matrix notation **B=SU**. The sources include both brain sources sensitive to sensory stimulation, and “noise” sources unrelated to stimulation. A role of data processing is to minimize the noise relative to the activity of interest.

### Acoustic data model

An auditory stimulus consists of a pressure waveform *p*(*t*), or *p_L_*(*t*) and *p_R_*(*t*) if the ears are independently stimulated, that is transduced and processed within the auditory system (cochlear filtering, haircell transduction, neural processing, etc.). To more easily relate the rapidly-varying sound signal to the slower brain signals measured by EEG, the pressure waveform *p*(*t*) must be transformed to a slower-varying representation *a*(*t*). Standard transforms (or “sound descriptors”) include the *temporal envelope* obtained by smoothing the instantaneous power *p*^2^(*t*) or the absolute value of the analytical signal obtained by the Hilbert transform, and the *spectrogram* obtained as a time series of short-term Fourier transform coefficients or of demodulated filterbank outputs mimicking the frequency selectivity found at multiple stages in the auditory system (“auditory spectrogram”’). For convenience, the acoustic representation is usually resampled to the same sampling rate as the EEG. It is well known that the same percept can arise from different stimuli (metamers) (Zaidi et al., 2013), and a yet larger set of stimuli may trigger EEG responses that are indistinguishable. A desirable feature of an EEG-relevant stimulus descriptor is to *discard* dimensions of the data that are not relevant to predict the response.

### Processing model

There are many linear processing techniques that may be used to enhance relevant components of EEG and acoustic representations. EEG sources responsive to sound may be enhanced in three ways: (1) by *spatial filtering*

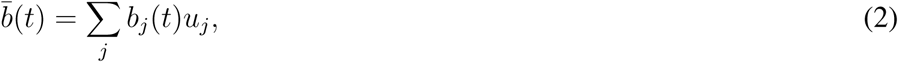

(2) by *spectral filtering* implemented as a finite impulse response (FIR) filter

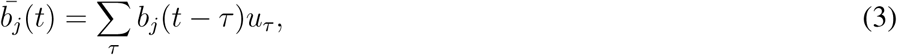

or (3) by a combination of the two implemented as a *multichannel FIR filter*

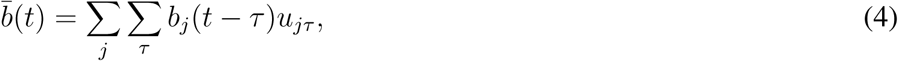

where *b̅*(*t*) is a linear combination of time-lagged brain signals. Spatial filtering allows unwanted sources to be suppressed based on the correlation structure between channels, whereas spectral filtering allows them to be attenuated based on their rate of fluctuation. The multichannel FIR filter of Eq. 4 subsumes both, as well as more complex "spatio-spectral" filters, for example for which different spectral filtering affects different spatial components.

The acoustic representation can likewise be enhanced with a FIR filter (for the stimulus envelope) or multichannel FIR filter (for a multichannel representation such as a spectrogram)

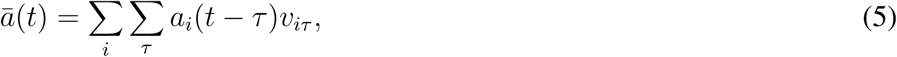

where *a̅*(*t*) is a linear combination of time-lagged audio descriptor signals. A FIR filter applied to the audio envelope or a channel of a spectrogram might select components based on their *modulation spectrum*, whereas the multichannel FIR applied to a spectrogram also captures more complex cross-spectral structure. These linear models allow great flexibility to optimize the relation between audio and EEG. However, the question remains as to how to find the appropriate coefficients *U_jτ_* and *v_iτ_* to apply to the EEG and audio signals respectively to maximize the correlation between the two.

### Canonical correlation analysis

Given two sets of multichannel data, CCA finds linear transforms of both that are maximally correlated. Given data matrices **X_1_** of size *T* × *J*_1_ and **X_2_** of size *T* × *J*_2_, CCA produces transform matrices **W_1_** and **W_2_** of sizes *J*_1_ × *J*_0_ and *J*_2_ × *J*_0_, where *J*_0_ is at most equal to the smaller of *J*_1_ and *J*_2_. The columns of **X_1_W_1_** are mutually uncorrelated, as are the columns of **X_2_W_2_**, while pairs of columns taken from both (“canonical correlate pairs”) are maximally correlated. The first pair of canonical correlates (CC) define the linear combinations of each data set with the highest possible correlation. The next pair of CCs are the most highly correlated combinations orthogonal to the first, and so-on. Assuming that **X_1_** and **X_2_** represent audio and EEG signals, possibly with time lags, CCA will produce weighted sums as in Eqs. 4, 5 that are maximally correlated, which is exactly what we are looking for.

### Dimensionality and overfitting

Reasoning in terms of vector spaces, the *J* EEG signals span a space of dimension *J* (at most) that contains all of their linear combinations. If *𝒯* time lags are applied to those signals (as in Eqs. 4, 5), the resulting space is of dimension *J𝒯* at most. Likewise, a spectrogram descriptor with *J′* bands spans a signal space of dimension of at most *J′*, or *J′𝒯′* if *𝒯′* time lags are applied. The weighted sums produced by CCA belong to these vector spaces. For each CC pair, the data-driven CCA process needs to find as many weights as the number of dimensions of the signal spaces (*J𝒯* + *J′𝒯* in this example). The number of dimensions is important because it determines the risk of *overfitting* in the data-driven calculation. Overfitting occurs when there are too many parameters relative to the amount of data and the algorithm latches on to spurious features.

There is a tradeoff between flexibility and overfitting: increasing the number of EEG channels and/or filter taps increases the ability to fit the data and resolve interesting sources and features, but this comes with a greater risk of misleading results.

Our strategy with respect to overfitting is twofold. First, the outcome of any analysis can be tested for overfitting by applying *cross-validation.* Second, overfitting itself can be limited by *dimensionality reduction* or *regularization* techniques, which are closely related (Jiang and Guo, 2007). Dimensionality reduction can be obtained by applying principal component analysis (PCA) and discarding principal components (PCs) with smallest variance. However this procedure implicitly equates variance to relevance, which may not always be appropriate (EEG artifacts and noise components often have high variance). Alternative approaches are considered in the Discussion.

### Filter bases

The processing model (Eq. 4 and 5) allows for FIR filters of order *𝒯* to equalize EEG and audio signals and attenuate irrelevant power. A larger *𝒯* captures temporal structure (of target and/or interference) on a longer-term scale, but there are more parameters and greater risk of overfitting. An alternative is to replace the set of time lags by a set of *𝒯* fixed filters with impulse responses of duration longer than *𝒯*, for example a logarithmic (or wavelet) filterbank. Again, it is useful to reason in terms of a vector space: *𝒯* time lags span the space of FIR filters of order *𝒯*, subspace of the space of all filters, whereas a *𝒯*-channel filterbank spans a different subspace with the same dimension, distinct from the first and including filters with different properties (in particular longer impulse responses). The number of parameters in either case is the same, but one subspace may better fit the requirements than the other. Tailoring the convolutional basis in this way is akin to *parameter tying* within a space of higher-order filters (LeCun et al., 2015). Whichever basis is chosen, CCA finds optimal coefficients on that basis.

### The quadratic trick

CCA can be made to handle nonlinear relations between data sets by replacing the data (or augmenting them) with a set of nonlinear transforms. A class of nonlinearity that deserves particular mention is *quadratic forms.* The rationale for considering it is the following. Suppose that there exists a source within the brain that displays a useful pattern of *power* (for example its power is correlated with the stimulus), but that source is weak and can only be observed after applying a spatial filter with coefficients *u_j_*. The values of these coefficients are unknown. However we note that the expression for the instantaneous power of the spatially filtered signal:

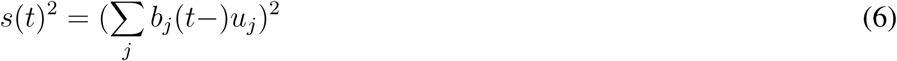

can be expanded as:

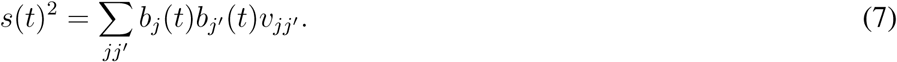

This expression is linear in the cross products *b_j_*(*t*)*b_j′_*(*t*), and thus we can apply linear techniques such as CCA to find a set of *v_jj′_* that maximize the correlation between the stimulus and the space of cross-products, from which we may derive an approximation of the optimal weights *u_j_* (de Cheveigné, 2012). This technique is equally applicable to spectral (FIR) filtering as to spatial filtering.

### Implementation

Processing is done in Matlab using routines from the Noise-Tools toolbox (http://audition.ens.fr/adc/NoiseTools/). Time lags and other transforms lead to large data matrices, and care must be given to computational constraints. The main ingredient used by the algorithms is the covariance matrix of the joint data set ([*X*_1_,*X*_2_]′*[*X*_1_,*X*_2_]) which can be calculated incrementally from subsets of the data, without loading all of the data into in memory. The main computational bottleneck is eigendecomposition of the covariance matrix (Matlab ‘eig’ function) with a cost that varies as *O*(*n*^3^) where *n* is the total number of columns.

### Evaluation

We implement several models, and evaluate each to determine the benefit of using CCA relative to other techniques. Model performance is quantified by calculating the correlation coefficient between stimulus-derived and EEG-derived quantities according to a cross-validation procedure. In brief, the model parameters are estimated using a subset of the data, and the correlation score is measured on the remaining data. This is repeated, leaving out different parts of the data in turn, and the final score is calculated as the mean of these estimates. When comparing models we take care to use the same number of parameters.

Auditory processing involves response latencies that are not perfectly known, and the convolutional transforms produced by CCA also contribute delays. In order to compensate for eventual differences in temporal alignment between stimulus and EEG, the entire process (model parameter estimation and correlation evaluation) is repeated for various time shifts *L* of the audio relative to the EEG. The optimal shift is determined from the peak of the correlation as a function of shift (see Figs. 1-3). This ensures an optimal temporal alignment for every model. This overall time shift *L* is distinct from the filter time lags *τ* that appear in some of the models (e.g. Eq. 4, 5).

**Figure 1:**
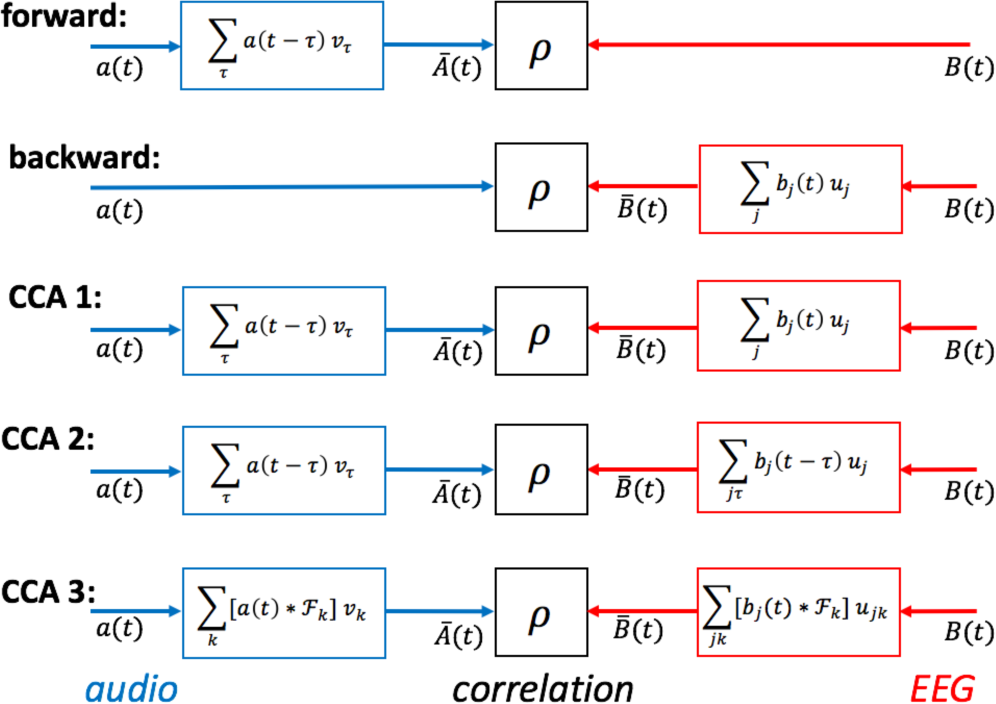
Overview of the models tested. In the forward model, the audio envelope is transformed and compared with the EEG. In the backward model, the EEG is transformed and compared with the audio envelope. In CCA model 1, a FIR filter is applied to the audio envelope and a spatial filter to the EEG. In CCA model 2, the spatial filter is replaced by a multichannel FIR filter. In CCA model 3, FIR filters are implemented using a bank of filters *𝓕_k_* instead of delays.

**Figure 2:**
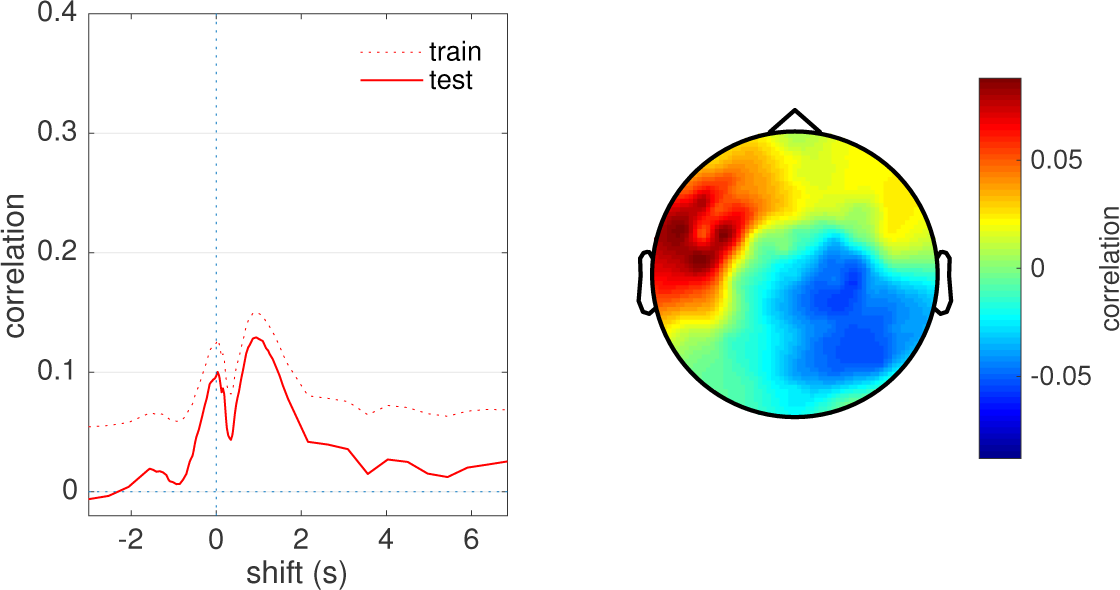
Backward model. Left: correlation score between the speech temporal envelope and the best linear combination of EEG, as a function of the overall temporal shift *L* of the stimulus relative to the EEG, for one subject (EL). A positive value of the abscissa means that the envelope has been delayed relative to the EEG. Dotted line is measured on the training set, full line on the test set. Right: spatial topography of the correlation between EEG and envelope at the optimal shift.

**Figure 3:**
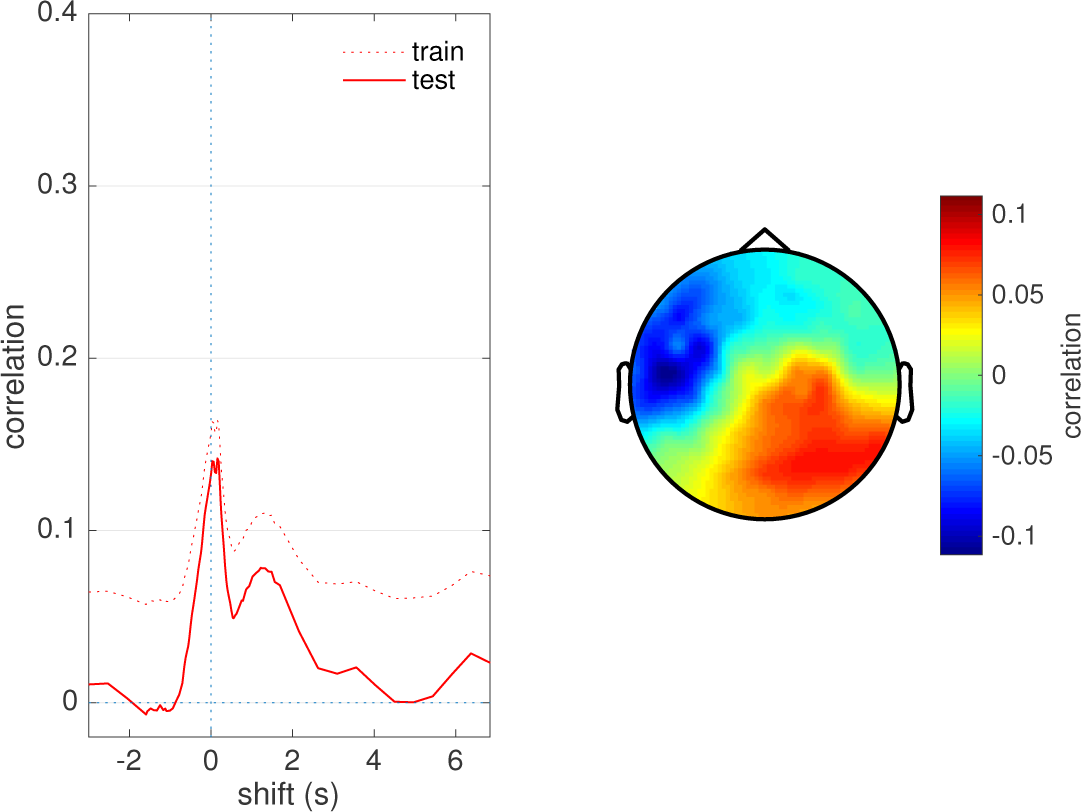
Forward model. Left: cross-validated correlation score between the FIR-filtered stimulus temporal envelope and the best EEG channel as a function of the overall temporal shift *L* of the stimulus relative to the EEG, for one subject (EL). Right: impulse response (top) and corresponding amplitude transfer function (bottom).

The decision to introduce a global time shift *L* distinct from *τ* was motivated by the desire to capture the effect of audio-to-EEG latency (fit by *L*) separately from that of spectral mismatch (fit by coefficients of *a*(*t* − *τ*) or *b*(*t* − *τ*), *τ* = 1… *𝒯*). The alternative of a wider range of *τ* to absorb latency would have entailed a larger number of parameters, and made it harder to make a fair comparison with models that don’t involve *τ* (e.g. Eq. 4). Introducing the parameter *L* allows each model to operate in optimal conditions.

Models are also evaluated by designing a *classifier* for a simple match-vs-mismatch classification task that involves deciding which of two segments of audio gave rise to a particular segment of EEG data of duration *D.* Segment duration is varied as a parameter, shorter durations being harder to classify because they contain less discriminative information. Again, cross-validation is used to control for overfitting: the classifier is trained on a subset of the data, and the classification score measured on unseen data. The classifier is trained and tested using the time shift *L* that maximized correlation.

### Evaluation data

The algorithms are evaluated using a database of EEG responses to natural speech reported in a previous study (Di Liberto et al., 2015). Full details of the stimulus and recording conditions are given in that reference. In brief, EEG data were recorded from 8 subjects using a 128-channel Biosemi system with standard electrode layout, at 512 Hz sampling rate (data from 2 additional subjects recorded with a 160-channel system were not used). Each subject listened to 32 speech excerpts, each of duration approximately 155s from an audio book presented diotically via headphones, for a total of approximately 1.4 hours. The EEG data were downsampled to 64 Hz and detrended by subtracting a 10th-order polynomial fit weighted to exclude outliers using a robust detrending routine (de Cheveigné and Arzounian, in review). The STAR algorithm (de Cheveigné, 2016) was used to suppress channel-specific noise, and the data were convolved with a boxcar window of duration 20 ms to suppress 50 Hz and harmonics. The data were then high-pass filtered with a Butterworth filter of order 2 and cutoff 0.1 Hz and rereferenced to the mean over channels. To calculate the stimulus’ temporal envelope, the stimulus (sampled at 44100 Hz) was squared, smoothed by convolution with a square window of width 15.6 ms (1/64 Hz), downsampled to 64 Hz, and then raised to the power 1/3.

### Models tested

To allow comparisons, we implemented backward and forward models, and three versions of the CCA model as schematized in Fig. 1.

In the *backward model* the stimulus representation *a*(*t*) is inferred or “reconstructed” as a transform *b̅*(*t*) of the EEG, for example a spatial filter:

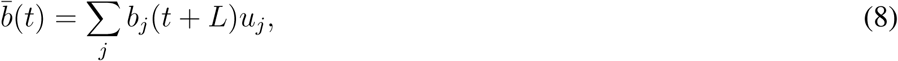

or spatiotemporal filter:

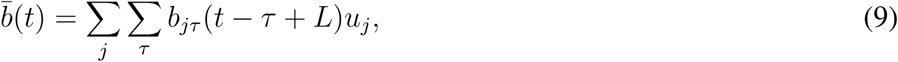

where the weights *u_j_* are calculated as the cross-correlation between the audio and time-lagged EEG. This model was simulated based on Eq. 8 (spatial filter). The 128 EEG channels were first submitted to PCA and the first 80 PCs retained, so the model had 80 parameters. The overall time shift *L* was varied over a range from -3 to +7 s, to find the highest correlation coefficient between *a*(*t*) and *b̅*(*t*). A variant of the model was simulated based on Eq. 8 (spatiotemporal filter) with 17 time lags, again with PCA to reduce the number of parameters to 80 (see Results).

In the *forward model* the response of a brain signal channel *b_j_*(*t*) is predicted from a transform *a̅*(*t*) of the stimulus, for example a weighted sum of time-lagged stimulus envelope samples (FIR filter):

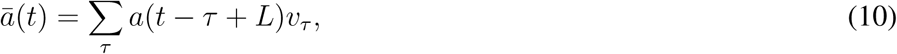

where the time shift *L* absorbs any temporal misalignment between stimulus and EEG. To give this model the best chances to succeed, the prediction was applied to the optimal linear combination of channels found by the backward model, rather than to a particular brain signal channel. The model was applied with 80 values of the delay *τ* (spanning 0 to 1.25 s), so this model too has 80 parameters.

In *CCA model 1*, both the stimulus and the EEG are transformed according to:

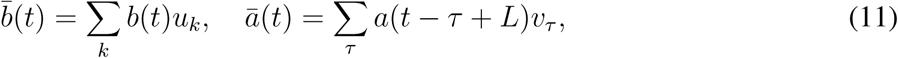

where the shift *L* is varied to absorb any mismatch in response or processing latency. CCA was applied to a set of 40 time lagged envelope signals and a set of 40 principal components of the EEG, producing 40 canonical correlate pairs. This model too has 80 parameters.

In *CCA model 2*, time lags are applied also to the EEG channels:

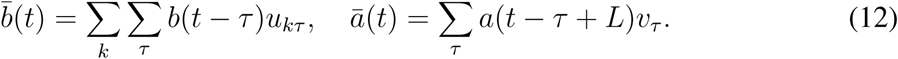

The EEG was submitted to PCA and the first 60 PCs were selected and submitted to 10 time lags. The compound matrix (600 channels) was submitted to PCA and 40 PCs retained, while the stimulus envelope was split into 40 time-lagged versions. This model too has 80 parameters. In a variant of this model (model 2+), 80 PCs were retained for both EEG and audio, so the model had 160 parameters.

In *CCA model 3*, time lags are replaced by a filter bank, motivated as follows. The stimulus envelope and EEG both include long-term fluctuations on the order of seconds. For example, these could reflect slow patterns shared by stimulus and response (e.g. phrase or semantic structure), or else slow artifacts that need to be removed. To resolve them requires filters with a long impulse response which, if implemented as a FIR, would require a filter of high order and thus many parameters leading to overfitting. For example, at 64 Hz sampling rate a two-second impulse response requires 128 taps, which applied separately to 50 EEG PCs (multichannel FIR) involves 6400 parameters. To address this issue, one can use a set of filterbank output channels as a “basis” for the space of filters, instead of time lags. For a *𝒯*-channel filterbank the number of parameters is the same as for *𝒯* time lags, but the filter subspace spanned is different. With a logarithmic or wavelet filterbank the basis can include both narrow filters with long impulse response, and wider filters with shorter impulse response. CCA model 3 was tested with a dyadic bank of FIR bandpass filters with characteristics (center frequency, bandwidth, duration of impulse response) approximately uniformly distributed on a logarithmic scale. There was a total of 21 channels with impulse response durations ranging from 2 to 128 samples (2 seconds). The filterbank was applied to the stimulus envelope, yielding 21 filtered signals, and also to the EEG represented by 60 PCs, yielding 1260 filtered signals that were submitted to another PCA from which 139 PCs were retained. These two data sets (21 and 139 channels respectively) were then processed by CCA. Like the previous model (CCA model 2+), this model has 160 parameters. The numbers chosen for this and previous models are largely arbitrary, the aim being to put the models in a range where they perform reasonably well, and allow comparisons between models with the same number of parameters.

Match-vs-mismatch classification was performed using Linear Discriminant Analysis (LDA) trained on the model correlation scores. The ability of each model to identify matching EEG and audio segments was assessed using leave-one-out cross-validation. The classifier was trained on all time segments from 31 of the 32 trials and tested on all segments in the remaining trial. To examine classification accuracy as a function of test data duration, we performed the classification with time segments of duration 1-64 s. This provided between 4960 (for 1 s segments) and 64 (for 60 s segments) non-overlapping tokens for classification.

## 3 Results

This section compares the performance of the different approaches described in Sect. 2, and investigates how the multiple canonical correlate pairs revealed by CCA can be used for classification.

### 3.1 Comparison between models

In the paragraphs to follow, we compare forward and backward models with CCA, using correlation as a performance metric.

#### Backward model

Figure 2 (left, full line) shows the correlation coefficient between the spatially filtered EEG and the speech temporal envelope, with cross-validation, as a function of the shift *L*, for one subject. The correlation coefficient is large for values of *L* near zero and falls for larger positive or negative values. The dotted line shows the coefficient without cross-validation. Plots in all other figures represent cross-validated correlation coefficients. Figure 2 (right) shows the spatial distribution over the scalp of the correlation coefficient between stimulus envelope and individual EEG channels. Coefficients for individual channels approach ~0.09, whereas the linear combination (Eq. 8) yields a score of ~0.13, reflecting the benefit of the spatial filtering.

Other studies, e.g. (O’Sullivan et al., 2014), used a model similar to Eq. 9 in which the stimulus is modeled as a weighted sum of time-lagged EEG channels. We simulated such a model using a set of 17 time lags (spanning 0-250 ms) as described in the Methods, using PCA to reduce the number of parameters to 80 as in the previous model. The peak score with this version was 0.17, vs 0.13 with the previous model (Eq. 8), suggesting a benefit of reconstruction based on spatio-spectral filtering over purely spatial filtering.

#### Forward model

Figure 3 (left) shows the correlation coefficient between the filtered stimulus envelope *a̅_j_* (*t*) and the best EEG channel (the channel that yields the highest correlation score). As for the backward model, the correlation coefficient peaks for small values of *L* and falls for larger positive or negative values. Figure 3 (right) shows the impulse response (top) and transfer function (bottom) of the FIR filter for the best shift. The transfer function peaks near 1 Hz, suggesting that correlation is better at the lower end of the frequency range.

The backward model was calculated from 80 PCs, while the forward model involved 80 time lags, so the degrees of freedom were the same for both. The second model applies an FIR filter applied to the stimulus envelope to predict an EEG channel, whereas the second applies spatial filter applied to the EEG data to infer the stimulus envelope. Since these filtering operations address different sources of noise, it is worth considering combining the two. The next sections explore how to do so in a systematic way using CCA.

#### CCA model 1

This first model is designed to be most comparable to the forward and backward model. Subsequent models take advantage of the flexibility of CCA to introduce a series of improvements. Figure 4 (first panel, red) shows the correlation coefficients for the first canonical correlate (CC) pair as a function of time shift. Values are comparable to the forward and backward models, but CCA produces *multiple* CCs beyond the first (thin lines), several of which seem to show elevated correlation scores for a limited range of shifts *L*.

**Figure 4:**
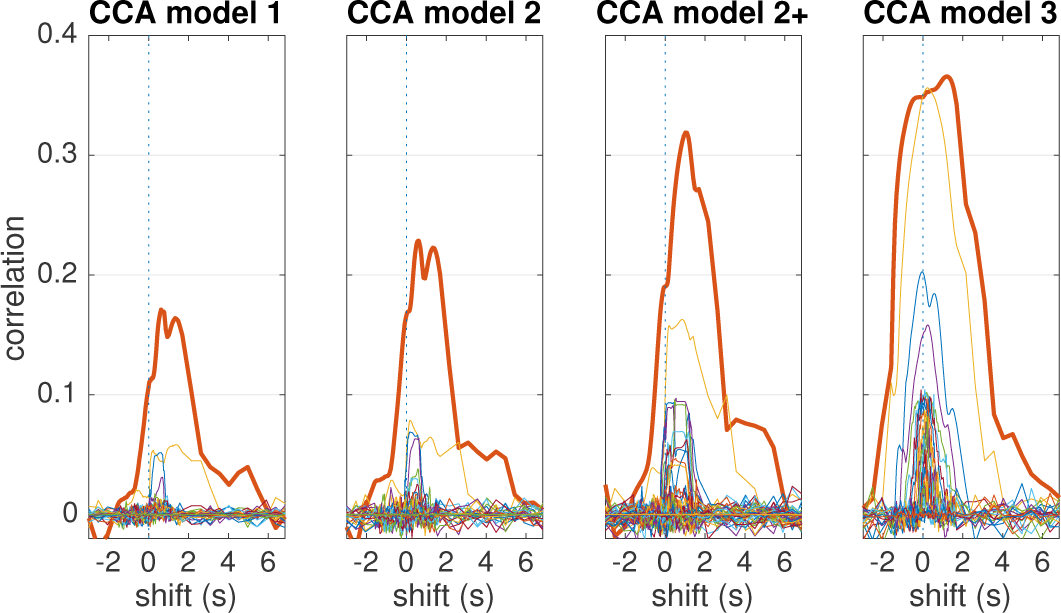
CCA models. See text for a description of each model. Each plot shows the correlation coefficient between canonical correlate pairs as a function of the temporal shift of the stimulus relative to the EEG, with cross-validation, for one subject (EL). One signal of each pair is derived by filtering the stimulus envelope with a FIR filter, and the other from the EEG by filtering with a spatial (leftmost) or multichannel FIR filter. The red line in each plot is the first (“best”) pair, other lines are for subsequent CC pairs.

#### CCA model 2

CCA model 2 allows time shifts for the EEG channels (Eq. 12). Figure 4 (second panel, red) shows shows the correlation coefficients for the first canonical correlate (CC) pair as a function of temporal alignment between stim-ulus and EEG. Values are higher than for the previous model, presumably as a result of the more flexible spatio-spectral filter. It is likely that the multichannel FIR filter adapts the spectral content of the EEG signals to that of the stimulus envelope. Thin lines show correlation coefficients for subsequent CC pairs.

Figure 4 (third panel, red) shows similar results assuming 80 time lagged stimulus envelope signals (instead of 40), and 80 PCs of time-lagged EEG signals (instead of 40) for a total of 160 parameters. Values are higher than for the smaller model, indicating that the more complex model gives a better fit to the data. This outcome was not a forgone conclusion, as the more complex model could instead have led to overfitting and thus a lower score with cross-validation. The larger correlation coefficients likely reflect the ability of higher-order FIRs to capture and resolve features on a longer time scale.

#### CCA model 3

Figure 4 (fourth panel, red) shows the correlation coefficient for the first CC pair as a function of time shift. The correlation coefficient approaches 0.4 at the best shift, indicating that CCA has discovered a transform of the stimulus that is highly predictive of a component extracted from the EEG by applying a spatio-spectral filter. In addition to this CC pair, there are additional well-correlated pairs, further discussed in Section 3.2.

The correlation coefficients for all subjects for these different models are summarized in Fig. 5 (the previous subject is the thick blue line). The values differ greatly between subjects, but the trends are overall similar. A Wilcoxon rank-sum test indicates that CCA conditions yield significantly larger scores than either forward or backward models (p<0.002, not corrected for multiple tests) and that CCA model 3 yields larger scores than any other model (p<0.002, not corrected).

**Figure 5:**
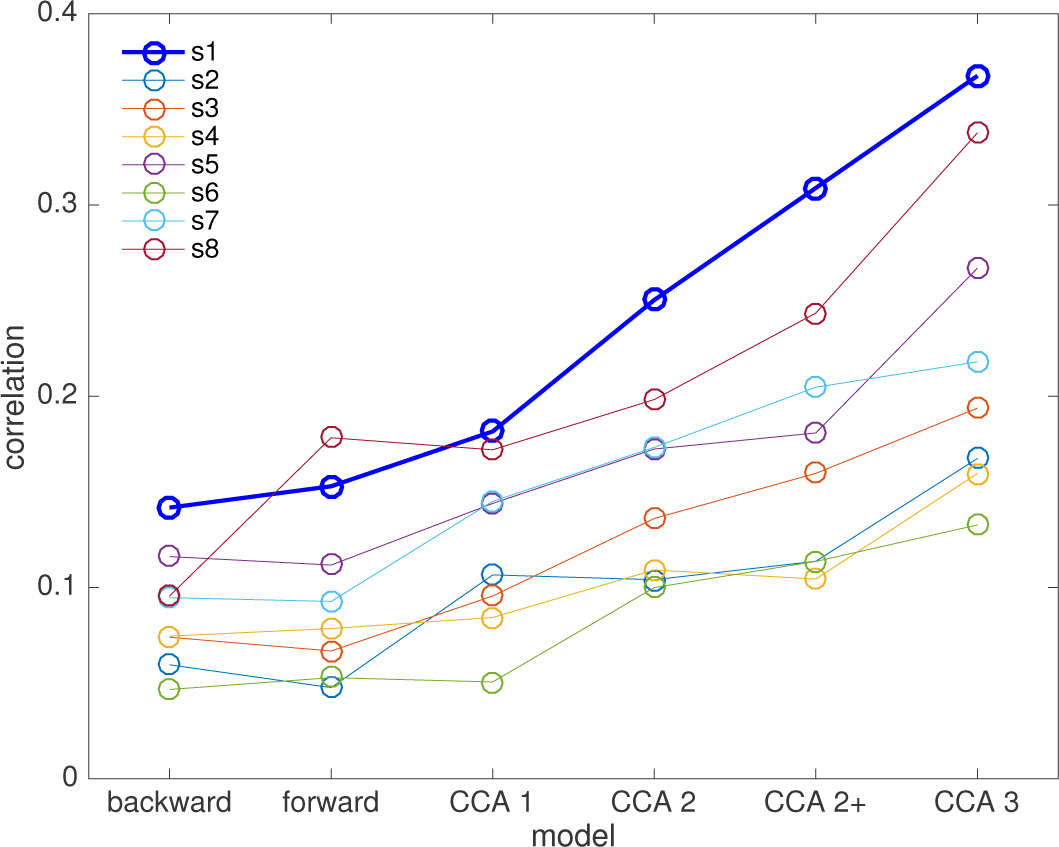
Best correlation scores for each subject as a function of the model. The thick blue line is the subject (EL) used for the comparison in Figs. 1-3 and Fig. 5

### 3.2 Exploiting multiple canonical components

A useful feature of CCA solutions is that there are *multiple* CCs with elevated correlation scores (Fig. 4). Each CC corresponds to a particular FIR-filtered stimulus-envelope signal, orthogonal to those of the other CCs, and that differs from the others in spectral magnitude and/or phase. In this sense, CCA performs a decomposition of the stimulus envelope in the modulation spectrum domain (Dau et al., 1997). Each CC is also associated with a spatial- or spatio-spectrally filtered EEG signal orthogonal to those of the other components, that differs from the others in spatial pattern and/or spectral magnitude and/or phase. CCA thus also decomposes the brain response into uncorrelated components, each mapped to a component of the stimulus envelope.

Figure 6 (top) shows the magnitude transfer functions of the FIR filters corresponding to the first 12 CCs of model 3 for subject EL. For most CCs the transfer function peaks in a particular spectral region, although the peak-to-skirt ratio is usually modest (roughly 4:1 in magnitude, 16:1 in power density). Each CC appears to cover a different spectral region but with considerable overlap between CCs. In this example, the transfer functions are peaked, and the peak frequencies roughly follow the order of the components, suggesting greater correlation values for lower frequencies, although the pattern is not perfect.

**Figure 6:**
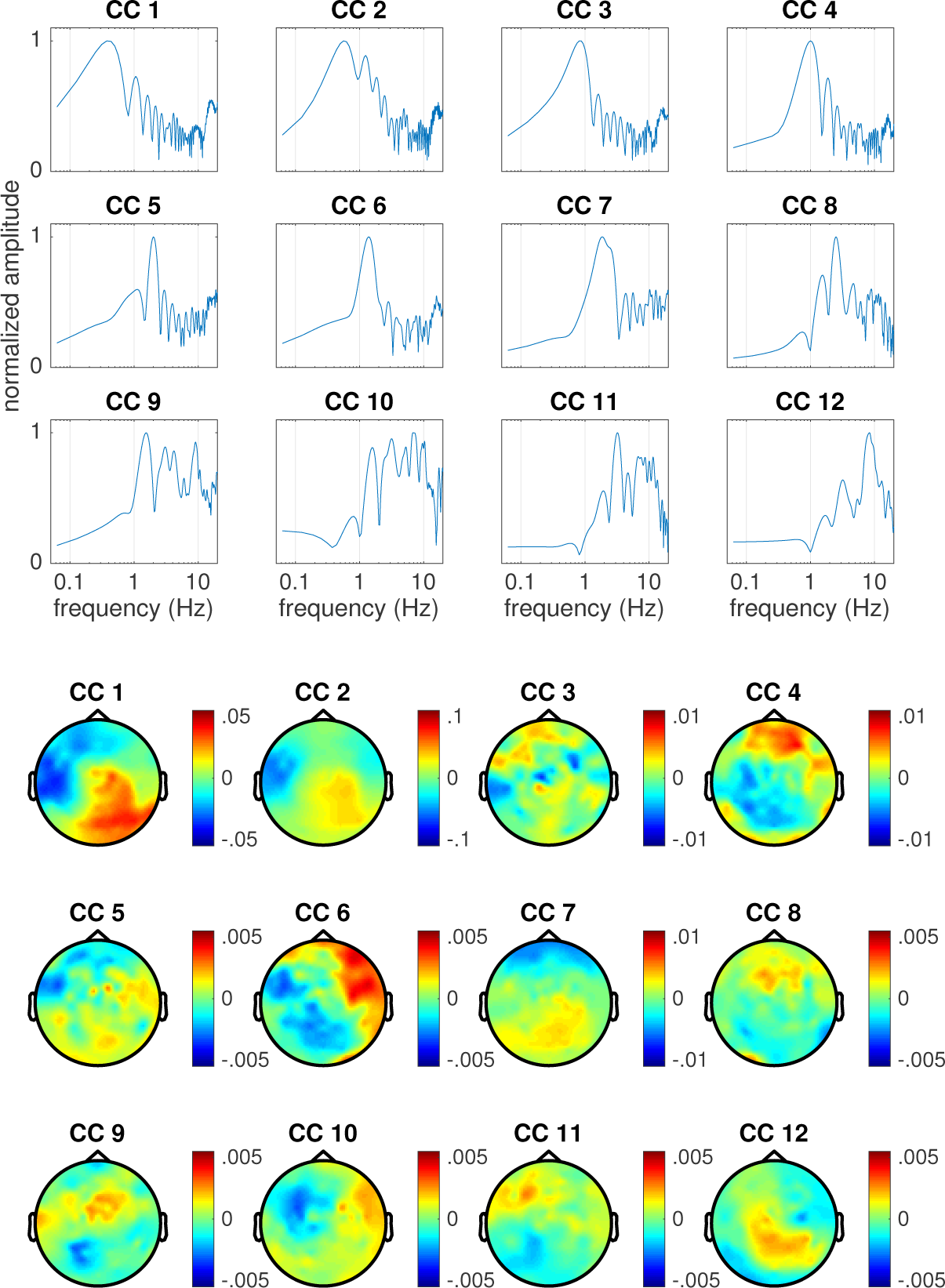
CCA model 3. Top: amplitude transfer functions of the first 12 CCA-derived FIR filters applied to the stimulus envelope normalized to a peak value of 1, for one subject (EL). Bottom: corresponding topographies. The value at each point of the topography is the normalized cross-correlation coefficient between the EEG component waveform and the EEG channel waveform. Signs are arbitrary. Note that the color scale differs between components, reflecting the lower SNR of higher-order components.

Figure 6 (bottom) shows the corresponding EEG topographies, calculated as the cross-correlation between the FIR-filtered envelope signal and the raw EEG channels (the polarity of each component is arbitrary). The topography gives a rough indication of the spatial substrate of the EEG component of the CC pair. Whereas the time-courses of activation are mutually orthogonal, there is no such constraint on the associated topographies, and indeed several seem quite similar. The patterns for weaker components tend to be noisy (note the different color scales for each CC).

As argued elsewhere (de Cheveigné and Parra, 2014) it is not possible to establish a one-to-one correspondence between components and specific sources within the brain. Component waveforms are mutually uncorrelated, whereas brain activity implicated in perceptual processing is likely to be correlated between sources. Even if the sources were uncorrelated, the source-to-component mixing matrix might not be diagonal. What can be said is that the components obtained from the analysis span a subspace of the EEG data containing the stimulus-related behavior. From a neuroscience perspective, the most interesting implication is that there are multiple processes, and that they cannot be subsumed by a single TRF model. CCA may be a useful tool to unravel them.

### 3.3 Classification

Another yardstick to measure a stimulus-response model is its performance in a classification task. Here we consider the task of deciding, given segments of stimulus and EEG of duration *T*, whether the EEG was in response to that stimulus or to an unrelated stimulus, using correlation scores as a feature as in prior studies. Classification involves determining whether a sample belongs to either of two distributions (Duda et al., 2012), in this case those of correlation scores for matched and mismatched segments respectively. Assuming Gaussian distributions with equal variance, the performance of a one dimensional linear classifier depends on the ratio of between-class to within-class variance, that can be quantified by the *d′* metric (distance between means divided by standard deviation). In the case of the forward and backward models, there is only one discriminant dimension, in the case of CCA there may be multiple discriminant dimensions.

Figure 7 (a) shows the *d′* scores for the first 6 CC pairs from CCA model 3, as a function of the size of the data segment over which the correlation between pairs is calculated. The *d′* values increase as the data segment becomes longer (as there is more information and estimates are more stable), and are greater for the first than for subsequent CC pairs. Values for the first few pairs are also greater than for the forward and backward models (Fig. 7 (c)). CC pairs that lead to large correlation scores tend to show large values of *d′* (Fig. 7 (b)), although the ratio between *d′* and correlation tends to be smaller for pairs for which the component waveforms are dominated by low frequencies (coded as blue) rather than higher frequencies (coded as red). Frequency content is quantified here by calculating the centroid of the power spectrum of the component waveform.

**Figure 7:**
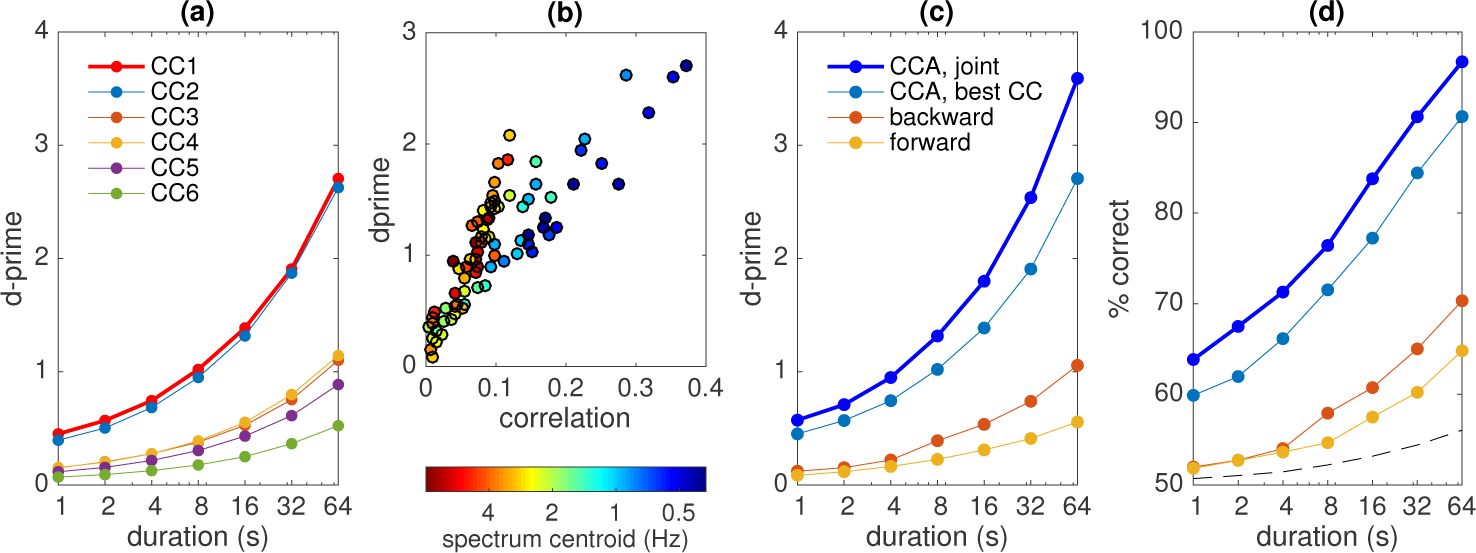
Match-vs-mismatch classification. (a) *d′* metric for the correlations of the first 6 CC pairs from CCA model 3, calculated over intervals of various durations. (b) Scatterplot of *d′* values (for an interval of 64s) versus correlation for all subjects and CC pairs up to rank 12. The color indicates the spectral centroid (in Hz) of the audio envelope-derived component of the pair. (c) *d′* metric for the forward and backward models, the first CC pair of CCA model 3, and the joint multivariate LDA model (thick blue line). (d) Percentage correct scores for a linear discriminant classifier for the same models as in (c). The dashed line in Fig. 7 (d) shows the 95th percentile of the classification score attained for 1000 random permutations of the class labels.

Rather than using individual CCs, correlation values can be combined over CC pairs using multivariate Linear Discriminant Analysis (LDA), as illustrated in Fig. 7 (d) (thick blue line) for the principal discriminant dimension. As expected, combining features from multiple CC pairs leads to better discrimination than the best CC Fig. 7 (d) (thin blue line), itself better than either the forward and backward models (red and orange lines). The dashed line in Fig. 7 (d) shows the 95th percentile of the classification score attained for 1000 random permutations of the class labels. We chose a simple linear classifier for simplicity and ease of exposition. A companion paper describes results with more sophisticated classifiers and classification strategies.

The match-vs-mismatch task, also chosen for simplicity of exposition, differs from the cocktail-party task considered by other studies (which of two concurrent streams is attended by the listener), although it captures the main aspects of that task. Here, the classifiers were trained over a relatively large dataset (> one hour) and tested on segments of new data, analogous to what might occur in a practical system designed to control a device (see paragraph Applications of Sect. 4). Note that the durations indicated in Fig. 7 (d) do not take into account the impulse responses of filters implied in certain models, so the amount of data involved in each case may be longer. In other words, the nominal durations may not faithfully characterize the ability of a classifier to track rapid changes in attention. The issue of classification latency is complex (it depends on the time scales of attention-dependent features as well as noise) and is explored in a companion paper.

## 4 Discussion

This study used CCA to reveal a subspace of cortical activity well correlated with ongoing auditory stimulation (speech). CCA yielded more accurate predictions (larger correlation values) than simpler forward and backward models on the same data, and allowed better classification. The presence of multiple CC pairs with different spatial signatures suggests that the analysis taps a complex cortical process involving multiple sources. The multiple discriminative dimensions support a wider range of classification schemes, and relatively good classification scores were obtained with short data segments, which is a step towards the goal of controlling an external device (for example a hearing aid) on the basis of cortical signals measured by EEG.

This study builds upon a growing body of work exploring stimulus-response relations using system-identification techniques and decoding (Lalor et al., 2009; Lalor and Foxe, 2010; Power et al., 2011; Pasley et al., 2012; Power et al., 2012; Ding and Simon, 2012; Brandmeyer et al., 2013; Ding and Simon, 2013; Ding et al., 2014; O’Sullivan et al., 2014; Koskinen and Seppä, 2014; Treder et al., 2014; Di Liberto et al., 2015; Mirkovic et al., 2015; Baltzell et al., 2016; Crosse et al., 2016; Ki et al., 2016; O’Sullivan et al., 2017; Biesmans et al., 2017; Fiedler et al., 2017; Khalighinejad et al., 2017; Fuglsang et al., 2017). Invasive measurements such as ECoG (Mesgarani and Chang, 2012; Tankus et al., 2012; Zion Golumbic et al., 2013; Chan et al., 2014; Martin et al., 2014; Leonard and Chang, 2014; Mesgarani et al., 2014; Lotte et al., 2015; Martin et al., 2015; Herff et al., 2015; Rao et al., 2017) support reconstruction of detailed spectrotemporal or symbolic representations of the stimulus, while EEG and MEG have been related to coarser representations such as the stimulus waveform envelope, as used here. Whereas ECoG samples mass electric fields relatively close to their source, the quality of EEG and MEG signals is degraded by source-to-sensor mixing and additional sources of noise. Methods such as CCA can contribute to improve the quality of these signals.

### A complex stimulus-response relationship for continuous speech

Each CC pair is characterized by a *filter* applied to the speech envelope. Each such filtered envelope waveform is orthogonal to the others, and together they form an empirical decomposition that splits the stimulus envelope into components spectrally distinct in amplitude and/or phase (Fig. 6, top). This is analogous to decomposition over a *modulation filterbank,* a hypothesized component of perceptual processing (Dau et al., 1997; Ewert and Dau, 2000; Lorenzi et al., 2001; Singh and Theunissen, 2003; Joris et al., 2004; Ghitza, 2011; McDermott and Simoncelli, 2011; Giraud and Poeppel, 2012; Wang et al., 2012; Xiang et al., 2013) that has also been proposed as relevant for processing temporally modulated sounds such as speech (Elhilali et al., 2003; Sukittanon et al., 2004; Hermansky, 2010; Nemala et al., 2013). Early studies applied a bandpass filter to EEG data to optimize SNR, e.g. 2-35 Hz (Lalor and Foxe, 2010) or 2-8 Hz (O’Sullivan et al., 2014). Koskinen and Seppä (2014) applied a logarithmic filterbank (wavelet analysis) to MEG data, obtaining relatively high correlation values within each band, particularly for the lowest frequency bands. Crosse et al. (2015) performed a similar analysis on EEG data using a linear filterbank (bandwidth 2 Hz). Compared to a filterbank, the empirical decomposition provided by CCA allows filter parameters to be directly optimized in both amplitude and phase (at the risk of overfitting, see paragraph Caveats).

Each CC is also characterized by a *spatiotemporal filter* applied to the EEG data. Each filtered EEG waveform is orthogonal to the others, and together they form a decomposition of the cortical activity into components that are spectrally and/or spatially distinct. Evidence has been found for spatially differentiated responses as a function of selective attention (Power et al., 2012; O’Sullivan et al., 2014; Fuglsang et al., 2017), linguistic competence (Brandmeyer et al., 2013), intelligibility (Tiitinen et al., 2012), phonemic feature (Khalighinejad et al., 2017), or modulation frequency band (Wang et al., 2012; Koskinen and Seppä, 2014). The general pattern suggests an early response to acoustic features from primary regions, and later responses to higher-order features (e.g. phonetic) from multiple secondary cortical regions (Norman-Haignere et al., 2015). CCA captures this spatially distributed and temporally-staggered activity, although, as mentioned earlier, there is not a one-to-one correspondence between components and neural sources.

### Canonical Correlation Analysis

CCA is a powerful tool to find linear transforms to apply to time series of brain data (Correa et al., 2010a; Dähne et al., 2015). Like PCA it produces component time series that are mutually decorrelated, but whereas PCA maximizes variance, CCA is designed to maximize correlation with a second set of time series. CCA is symmetrical with respect to the datasets, and thus a similar transform is found for the second set that maximizes correlation with the first. Components come by pairs (canonical correlates), the first pair consisting of the linear combinations of either data set with the highest possible correlation between them.

CCA is intended for multichannel data, but can be applied to single channel data by augmenting them with time-lags. The solution (linear combination of time-lagged data) is then a *FIR filter*, the effect of which is to equalize the spectrum of the time series so as to maximize its correlation with the other dataset. Applying time lags to multichannel data results in *multichannel FIR filters* that exploit both spectral and spatial structure. The time lags can also be replaced by other convolutional transforms (such as the logarithmic filterbank we used here). Finally, applying nonlinear transforms to the data before CCA allows it to capture non-linear relations between data sets, that can also be found using deep learning techniques (Bießmann et al., 2009; Andrew et al., 2013; Dähne et al., 2015; Wang et al., 2015; Tang et al., 2017). CCA is a flexible tool to discover complex convolutional and non-linear relations between sets, but this flexibility is of course offset by the risk of overfitting entailed by the many parameters involved (see paragraph Caveats).

In a recent study CCA was used to characterize auditory and audio-visual EEG responses (Dmochowski et al., 2017). They applied time-lags to the stimulus representation but not both stimulus and EEG, and they did not consider a convolutional basis other than time lags, both of which provided significant improvements in our study. CCA was used by Biesmans et al. (2017) to combine information across channels of an auditory filterbank to maximize correlation with EEG, and by Koskinen and Seppä (2014) to maximize trial-to-trial reproducibility before relating stimuli to responses. CCA is one example of a wider class of methods that apply linear transforms to EEG to isolate components correlated between datasets (Bießmann et al., 2009; Dmochowski et al., 2012; Lankinen et al., 2014; Dmochowski et al., 2015; Sturm et al., 2015; Biesmans et al., 2017) or between modalities (Correa et al., 2010b; Sui et al., 2013).

### Caveats and cautions

As other data-driven analysis techniques, CCA is prone to overfitting and circularity (Kriegeskorte et al., 2009). The risk is aggravated because of the many parameters, particularly if multiple transforms of the data (convolutional and non-linear) are included. Overfitting may be detected and to some degree remediated using standard cross-validation and surrogate data techniques, and the tendency to overfit limited by dimensionality reduction, regularization, or prior constraints (Jiang and Guo, 2007; Lahti et al., 2009; Andrew et al., 2013). While dimensionality reduction and regularization are closely related, we prefer the former as it allows dimensionality to be reduced based on criteria other than that optimized by CCA (e.g. spectral content, or repeatability), and competing models can be put on a similar footing by giving them the same number of free parameters.

CCA, based on least-squares minimization, is prone to large-amplitude outliers and glitches that may dominate the solution. It may also pick up artifacts that happen to be correlated between data sets, such as power line interference, linear trends, or filtering transients. Introducing time lags (or filters) allows CCA to synthesize filters that isolate particular frequency regions, leading to higher correlation scores (compare CCA model 2 and 3 to CCA model 1). However, higher correlation might also arise merely from the greater serial correlation introduced by lowpass or narrowband filtering, that reduces the degrees of freedom within the data and thus exacerbates overfitting. Indeed, we observed that, for a given value of correlation, discriminability (*d′*) tends to be lower for components dominated by lower frequencies (Fig. 7 (b)).

Figure 7 (d) reports classification scores for data segments as short as 1 s, but it should be noted that these nominal durations do *not* include the length of the FIR filters. This should be taken into account when comparing models, or judging the ability of an algorithm to track fast changes in an application (see next paragraph).

### Relevance for applications

A motivation for this study was to develop a method to steer an auditory assistive device (e.g. hearing aid) using brain signals. Several studies have shown that it is possible to determine which sound source among several is the focus of attention using EEG (Power et al., 2012; O’Sullivan et al., 2014; Mirkovic et al., 2015; Fuglsang et al., 2017; Ki et al., 2016; Biesmans et al., 2017). Good performance has been reported for stimulus-response data of long duration (e.g. 30 or 60 s) for some subjects, but accuracy falls short of what would be needed for real-time control in a usable device. For example, a 95% accuracy (as reported by Biesmans et al. (2017) for their best subject at 30 s duration) would amount to one error in every 20 decisions. An error rate such as this is likely to be frustrating for a user, and having to wait tens of seconds for a decision is also not acceptable.

CCA brings our goal closer in at least three ways. The first is by allowing the representations of both stimulus and response to be optimized jointly, leading to higher correlation scores (Fig. 5) and better classification (Fig. 7). The second is by providing *multiple* discriminant measures that support multivariate classification as illustrated here with a simple multivariate LDA classifier (Fig. 7 (d)). The third is by providing a flexible framework that can harness additional streams of information (for example non-linear transforms of the data). The best scores reported here (Fig. 7) still fall short of what is needed for a useful device, but the CCA methodology opens many routes that may lead to better performance. In our opinion the value of CCA lies less in the scores reported here, however encouraging, as in the potential it offers for further development. A companion paper describes results with more sophisticated classifiers based on CCA.

The computational costs associated with CCA are mild. The solution requires eigendecomposition of a covariance matrix that can be calculated incrementally and updated in real time, and efficient standard methods are available for eigende composition. Once the solution has been derived, processing amounts merely to application of a multichannel FIR (Eqs. 4 and 5).

## Conclusion

CCA strips the brain response of variance unrelated to the stimulus, and the stimulus of variance that does not affect the brain response, and thus improves our observation of the relation between stimulus and response. The best performing CCA model produced multiple component pairs, each consisting of a FIR filter applied to the stimulus waveform, and a multichannel FIR filter (spatiotemporal filter) applied to the EEG signal. The spatial signatures of these filters were diverse, suggesting that the analysis is sensitive to multiple neural sources, although there is no simple mapping of components to sources. Each pair was also associated with a different FIR filter applied to the stimulus envelope, indicating sensitivity to different rates of change. Signal components isolated by CCA supported effective classification, in particular thanks to the fact that multiple discriminative components are available, bringing closer the perspective of a cognitively-controlled device based on recording of attention-sensitive correlates of auditory stimulation. CCA is of great interest as a tool to explore the effectiveness of transforms (e.g. nonlinear or convolutional) applied to the data, to further our understanding of perceptual processes and increase the effectiveness of applications.

## Acknowledgements

This work was supported by the EU H2020-ICT grant 644732 (COCOHA), and grants ANR-10-LABX-0087 IEC and ANR-10-IDEX-0001-02 PSL*, as well as a Ulysses grant (2015) from the French Ministry of Foreign Affairs and the Irish Research Council. It draws on work performed at the 2015 and 2016 Telluride Neuromorphic Engineering workshops. Giovanni Di Liberto was supported by an Irish Research Council Government (GOIPG, 2013-2017) of Ireland Postgraduate Scholarship.

